# SAMP: Identifying Antimicrobial Peptides by an Ensemble Learning Model Based on Proportionalized Split Amino Acid Composition

**DOI:** 10.1101/2024.04.25.590553

**Authors:** Junxi Feng, Mengtao Sun, Cong Liu, Weiwei Zhang, Changmou Xu, Jieqiong Wang, Guangshun Wang, Shibiao Wan

## Abstract

It is projected that 10 million deaths could be attributed to drug-resistant bacteria infections in 2050. To address this concern, identifying new-generation antibiotics is an effective way. Antimicrobial peptides (AMPs), a class of innate immune effectors, have received significant attention for their capacity to eliminate drug-resistant pathogens, including viruses, bacteria, and fungi. Recent years have witnessed widespread applications of computational methods especially machine learning (ML) and deep learning (DL) for discovering AMPs. However, existing methods only use features including compositional, physiochemical, and structural properties of peptides, which cannot fully capture sequence information from AMPs. Here, we present SAMP, an ensemble random projection (RP) based computational model that leverages a new type of features called Proportionalized Split Amino Acid Composition (PSAAC) in addition to conventional sequence-based features for AMP prediction. With this new feature set, SAMP captures the residue patterns like sorting signals at around both the N-terminus and the C-terminus, while also retaining the sequence order information from the middle peptide fragments. Benchmarking tests on different balanced and imbalanced datasets demonstrate that SAMP consistently outperforms existing state-of-the-art methods, such as iAMPpred and AMPScanner V2, in terms of accuracy, MCC, G-measure and F1-score. In addition, by leveraging an ensemble RP architecture, SAMP is scalable to processing large-scale AMP identification with further performance improvement, compared to those models without RP. To facilitate the use of SAMP, we have developed a Python package freely available at https://github.com/wan-mlab/SAMP.

## Introduction

Between the period of 1940s and 1960s, one of the greatest breakthroughs was the development of antibiotics [1], a remarkable medication that has saved thousands of lives by defeating various infectious diseases [2–6]. However, the long-term and rapid increase of antibiotic use for disease treatment in large populations has resulted in the emergence of drug resistance in pathogens [7–11]. The Centers for Disease Control and Prevention (CDC) has reported that drug-resistant bacteria caused around 2.8 million infections and more than 35000 deaths in the United States [12]. According to the World Health Organization (WHO), approximately 700,000 patients worldwide die from drug-resistant bacterial infections every year, and the total number of deaths is predicted to increase to 10 million by 2050 [13], making it an urgent challenge in the healthcare system [14,15]. Therefore, expanding a large range of new antimicrobial agents to fight against pathogens is essential to relieve the huge burden of global health [16].

Antimicrobial peptides (AMPs) are amino-acid-based oligomers or polymers, naturally widespread in all forms of life, such as bacteria, animals, and plants [17–19]. They have played an important role in protecting organisms from infectious diseases for millions of years, serving as the first line of defense against pathogens through interrupting pathogen-associated molecular processes in the innate immune system [20–24]. It has been suggested as excellent candidates for developing a new generation of antibiotics due to their special ability to kill multi-resistant microorganisms, including bacteria, fungi, parasites, and viruses [25–29]. In addition, substantial evidence indicates that AMPs can recruit immature phagocytic and dendritic cells, leading to the destruction of cancer cells and healing wounds in certain areas [24,30]. Cationic and hydrophobic residues are two main characteristics of linear AMPs, enabling sequences to fold into amphipathic secondary structures which tend to disrupt negatively charged membranes of pathogens while sparing the healthy eukaryotic cells. Moreover, the enrichment of cholesterol and neutral phospholipids in membranes also makes eukaryotic cells much less susceptible to AMPs [31,32]. The main mechanism of AMPs’ action is to form pores and micellization in cell membranes or directly cause osmotic shock when present in high concentration [33,34]. Additionally, binding to specific cytosolic macromolecules is another way of AMPs to inhibit the synthesis process of the cell wall or ribosomes [35,36]. Hence, AMPs can interact with many different components of bacteria for multiple hits, while traditional antibiotics are typically designed to target one specific enzyme. The broad interactions of AMPs make it difficult for bacteria to develop resistance in a short time [37–39].

Natural AMPs discovery typically relies on traditional time-consuming and labor-intensive wet experiments, resulting in low efficiency. Therefore, to find natural AMPs in a more efficient and convenient way, it is necessary to develop in-silico predictive models to identify possible AMP candidates prior to synthesis and wet lab testing. In the past decade, numerous computational models based on various algorithms, such as support vector machine (SVM) [40], random forest (RF) [41] and logistic regression (LR) [42], have been introduced to identify peptides [43]. Most recently, Huang et al. [44] constructed a sequential model ensemble pipeline (SMEP) consisting of multiple steps, including empirical selection, classification, ranking, regression, and wet-lab validation. Algorithms, like boosting method (XGBoost) [45], RF as well as deep learning such as the convolutional neural network (CNN) [46] and the long short-term memory (LSTM) [47], were applied in different modules. With SMEP, a series of potent AMPs from the entire search space of peptide libraries were identified accurately within a short period of time. In another study [48], multiple natural language processing neural network models, including LSTM layer, Attention layer and Encoder Representations from Transformers (BERT) [49], were combined to form a unified pipeline which has been used to mine functional peptides from metagenome data for in-depth investigations. Based on the algorithms applied in the prediction models, they can be divided into two main categories. Models in the first category are based on the deep learning (DL) architectures, like AMPScanner V2 [50] and Deep-AmPEP30 [51]. AMPScanner V2 applied deep neural networks (DNN) [52] with convolutional, maximal pooling and LSTM layers for AMPs prediction. Deep-AmPEP30 based on convolutional neural network (CNN) with two convolutional layers, two maximum pooling layers, and one fully connected hidden layer to identify specifically short length AMPs which contain fewer than 20 amino acids. As for the second category of models, conventional machine learning (ML) algorithms are generally exploited, such as iAMPpred [53] which used SVM to classify positive or negative peptides. Previous studies [54] indicated that DL models did not always outperform conventional ML models due to the modeling complexities and/or modeling overfitting during the process of DL model construction based on training limited AMPs. Therefore, DL models are not necessarily the most suitable approach for AMPs identification [54]. Nonetheless, no matter the ML or DL based methods, existing computational methods rely primarily on features derived from the composition, physicochemical and structural features of the peptide sequence. These features may not be sufficient to fully express the rich information contained in antimicrobial peptides and there is still considerable room for enhancing accuracy of AMP prediction.

To address the aforementioned concerns, we propose herein an ensemble random projection (RP) [55] based computational model named SAMP, for which we develop a new type of features called proportionalized split amino acid composition (PSAAC) [56] in addition to conventional sequence-based features to improve the prediction performance of AMP identification. Residue patterns such as sorting signals at around both the N-terminus and the C-terminus could be captured by SAMP by using this enhanced feature set, while also remaining the sequence order information extracted from the middle region fragments. Meanwhile, we demonstrate that SAMP outperforms existing state-of-the-art methods in terms of accuracy, Matthews correlation coefficient (MCC), the geometric mean of recall and precision (G-measure) and F1-score, including iAMPpred and AMPScanner V2, when benchmarking on both balanced and imbalanced datasets from different natural peptide groups, including humans, bacteria, amphibians and plants. Furthermore, we integrate an ensemble RP architecture into SAMP to strengthen the ability of handling large-scale AMP screening while achieving enhanced performance compared to those without RP. We believe SAMP will play a significant role in AMP identification, complementary to existing AMP identification approaches.

## Materials and Methods

### Datasets

The positive data set for natural AMPs was accumulated in the antimicrobial peptide database in the past 20 years [57,58]. The negative data set was extracted from UniProt by excluding peptides/proteins annotated with key words such as “antimicrobial”, “antibacterial”, “antiviral”, and “antifungal” [59]. To benchmark the performance of SAMP and other state-of-the-art approaches, we selected two sets of training data reported in the literature. As many existing approaches only provide web servers which have already been trained in different training data, to make a fair comparison, we will compare SAMP with those approaches based on the same training dataset based on which the corresponding web servers were trained. Specifically, the first set consists of 984 positive and 984 negative antimicrobial peptide sequences obtained from [53]. This set is used to train our model and compare our proposed model SAMP with iAMPpred (**Fig. 1A**). The second set consists of 2021 positive and 2021 negative antimicrobial peptide sequences from [50], as shown in **Fig. 1B**. This set is used to train our model and compare SAMP with AMPScanner V2 [50].

**Fig. 1.**
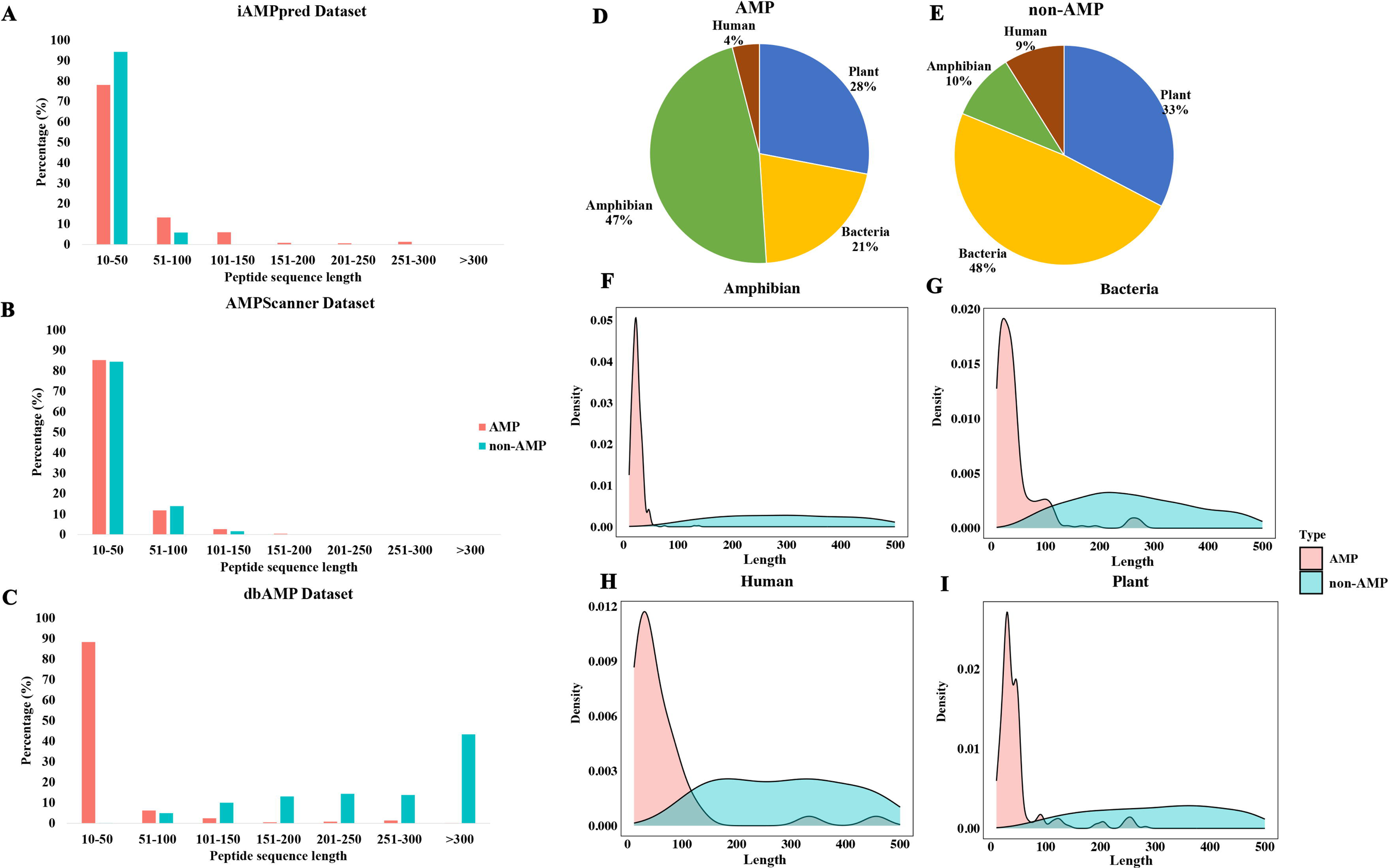
Peptide sequence distribution of AMPs and non-AMPs in benchmarking datasets. (**A-C**) The peptide sequence distribution of AMPs and non-AMPs collected from the iAMPpred dataset **(A**), the AMPScanner V2 dataset **(B)** and the dbAMP dataset **(C**), respectively. **(D-E)** Species breakdowns of AMPs **(D)** and non-AMPs **(E)** in the dbAMP dataset. **(F-I)** Species-specific peptide sequence distribution of AMPs and non-AMPs in the dbAMP dataset, including amphibian **(F**), bacteria **(G**), human **(H)** and plant **(I)**.

In addition, independent testing data were collected from the dbAMP database [60], containing AMP and non-AMP sequences (**Fig. 1C**). Specifically, we chose the AMP and non-AMP datasets across four different species: plants, bacteria, amphibians, and humans, which were originally collected in the APD [57,58] database. Given the varying peptide sequence length distributions of our AMP datasets (**Figs. 1A-C**), we filtered out sequences shorter than 10 amino acids and longer than 500 amino acids. The sequences containing non-standard amino acids were also removed. In the dbAMP benchmark dataset (**Fig. 1D-E**), in total, there are 1089 AMPs and 9732 non-AMPs. As found in the APD database, amphibians predominantly constitute the natural AMP sequences, while bacteria form the majority in the non-AMP sequences. Specifically, for the AMPs (**Fig. 1D**) of the dbAMP dataset, around half are amphibian, one third belong to plant, and one fifth are bacteria. On the contrary, in the non-AMP cases (**Fig. 1D**), amphibian sequences account for only 10%, and half of them are bacteria. Interestingly, human sequences constitute less than 10% in both AMPs and non-AMPs (**Figs. 1D-E**). While for the amino acid sequence length distribution (**Figs. 1F-I**), most AMPs for all species are with shorter amino acid sequences compared to non-AMPs, suggesting significantly different sequence distributions between AMPs and non-AMPs. However, it is unlikely to use the length of peptide sequences to determine whether a peptide is an AMP or non-AMP, given that a significant portion of AMPs are also overlapped with non-AMPs, especially for bacteria, human, and plant (**Figs. 1F-I**).

### Feature extraction

#### Conventional features

First, we embedded the string of peptide sequences into categories of numeric feature vectors similar to those proposed in [53], which includes amino acid sequence compositional features and physio-chemical (PHYC) features. The compositional features include amino acid composition (AAC), pseudo amino acid composition (PAAC), and normalized amino acid composition (NAAC). The PHYC features consider the hydrophobicity, net-charge, and iso-electric point of peptide sequences, and were calculated using the ‘Peptide’ package [61] in R.

#### Proportionalized split amino acid composition (PSAAC)

In addition, to maximally extract peptide sequence information, we propose a new compositional feature called proportionalized split amino acid composition (PSAAC). This concept refines the split amino acid composition (SAAC) approach, which differentiates between the amino acid compositions at the N and C-terminus of protein sequences [62,63]. PSAAC adapts this concept specifically for peptide sequences, dividing them into distinct segments according to proportions defined by the users. Given a peptide sequence 𝒫 of length *L*, we split it into 3 segments using proportions (or percentage), *p*_1_, *p*_2_ and *p*_3_, where *p*_1_, *p*_2_ and *p*_3_ represent the proportion of amino acid segments for the N-terminus region, the middle region and the C-terminus region, respectively, and *p*_1_ + *p*_2_ + *p*_3_ = 1. The lengths of these segments, *L*_1_, *L*_2_, and *L*_3_, are:

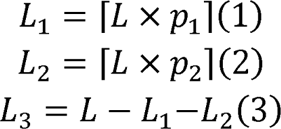

The segments are:

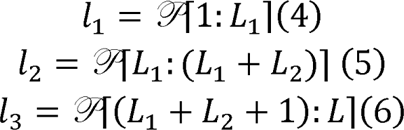

Now, let 𝔄 be the set of 20 standard amino acids. The amino acid composition (*AAC*) in segment *1_i_* for *X* ∈ 𝔄 is given by:

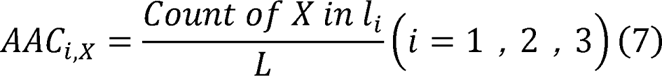

Note that the count of the *X* in each segment was divided by the whole length of the peptide sequence. Then, the proportionalized split amino acid composition (PSAAC) is:

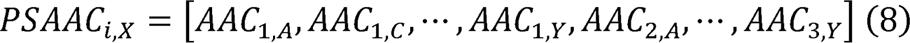

Previous studies [64,65] have reported that some sorting signals exist in the short segments of amino acid sequences around the N-terminus, representing special information of amino acid composition. In other words, different regions of a protein sequence can provide extra information. For example, some specific regions may form structural domains that determine the function of proteins, such as binding sites for other molecules, active sites for enzymes, or domains for protein-protein interaction [65]. The PSAAC feature captures the residue patterns around both the N-terminus region and the C-terminus region, while also retaining the sequence order information from the middle region. Based on peptide sequences from [50], the amino acid compositions for each segment (e.g., the N-terminus region, the C-terminus region, and the middle region) were calculated respectively. Non-standard amino acid residues are ignored. As shown in **Fig. 2**, Leucine and Glycine are the most abundant amino acids in AMPs as found originally in the APD, and non-AMPs dataset respectively (**Figs. 2A-B**). There are obvious differences in the composition of each amino acid at the N-terminus, the C-terminus and middle region for both datasets (**Figs. 2C-D**). In the non-AMPs dataset, the least abundant amino acids are Tyrosine, Methionine and Tryptophan at the N-terminus, middle region, and the C-terminus respectively. Conversely, Leucine is the most abundant in all three segments. For the AMPs dataset, Glycine is the most abundant at both the N-terminus and the middle region, while Lysine is the most abundant at the C-terminus. The amino acids with the lowest content in the AMPs dataset are Histidine at the N-terminus, Methionine in the middle region, and Tryptophan at the C-terminus. Then, all the features are scaled by subtracting the mean from each column and dividing it by the standard deviation. For the data collected from AMPScanner V2 and dbAMP, the amino acid distribution at the N-terminus, the C-terminus and middle region is also investigated and shown in **Supplementary Figs. S1**-**S2**. For each peptide sequence, the PSAAC feature will be generated with 60 dimensions. We note that *p*_1_, *p*_2_ and *p*_3_ are user-defined hyperparameters that allow flexible sequence context extraction and the number of splits can also be customized, and we used three splits because of its prominent performance.

**Fig. 2.**
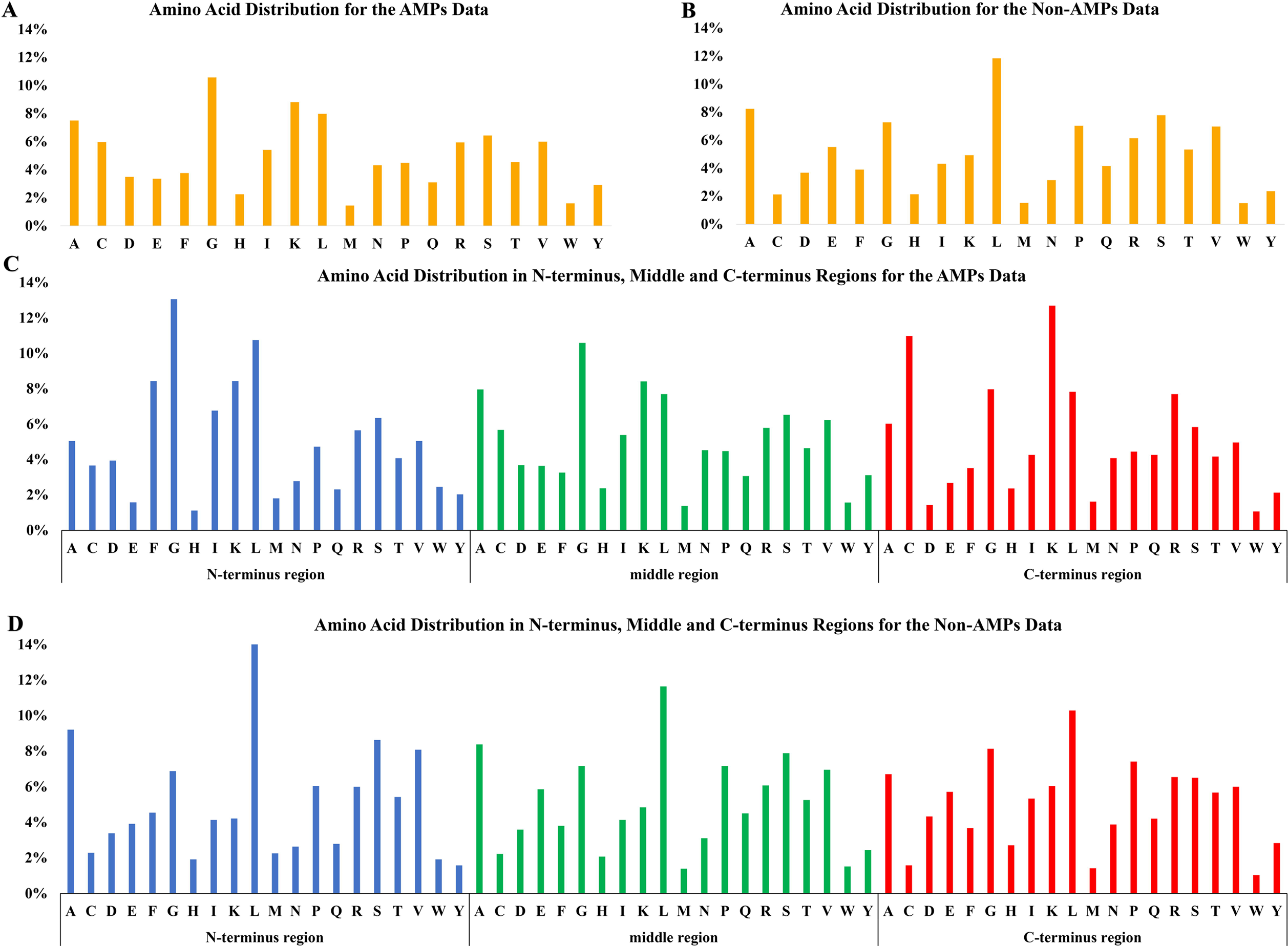
Amino acid distribution in AMPs and non-AMPs datasets based on the dataset collected from iAMPpred. Amino acid distribution of all sequences in (**A**) AMPs and (**B**) non-AMPs dataset. Distribution of amino acid sequences in the N-terminus region, the middle region and the C-terminus region of (**C**) AMPs Dataset and (**D**) non-AMPs dataset.

### Random projection

Random projection (RP) is a dimension reduction technique proposed based on the Johnson-Lindenstrauss lemma [66]. For our experimental analysis, we used the Gaussian random matrix as our random projection matrix, which is generated from the following the distribution 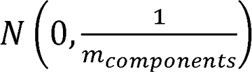 where *m_components_* represents the number of dimensions to which the data is to be reduced. In cases where the number of feature dimensions is high, the use of random projection can greatly speed up the model training process. In our experiments, the optimal number of components to be kept was determined by the model training step using a grid-search approach. We also enabled the option of using a sparse matrix as the random projection matrix in our package.

For dimension reduction, from original *R* dimension to the reduced *r* dimension, a very sparse random matric **Q** ∈ ℝ^r×R^ is designed to reduce the computational complexity [67]. Specifically, elements of Q (i.e., *q_i,j_*) are defined as:

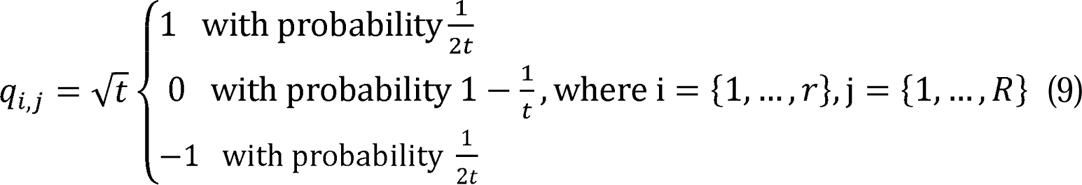

As suggested by [67], we select *t*=√R.

### Ensemble learning

We use an ensemble learning model in SAMP (**Fig. 3**) where given the training and testing feature matrices **M***_train_* and **M***_test_* that have been scaled, and whose scaling process will be detailed in the feature scaling session, we first applied random projection on the matrices respectively to get the new feature matrices **M***\*_train_* and **M***\*_test_* in a lower dimension. We then used the SVM as our base model to train and test on **M***\*_train_* and **M***\*_test_* respectively. The decision function scores on the testing data are recorded. We repeated the above steps for 10 times to stabilize the result of random projection, where randomness is often introduced when generating the random projection matrix. Finally, the decision function scores in each iteration are averaged to get the final scores.

**Fig. 3.**
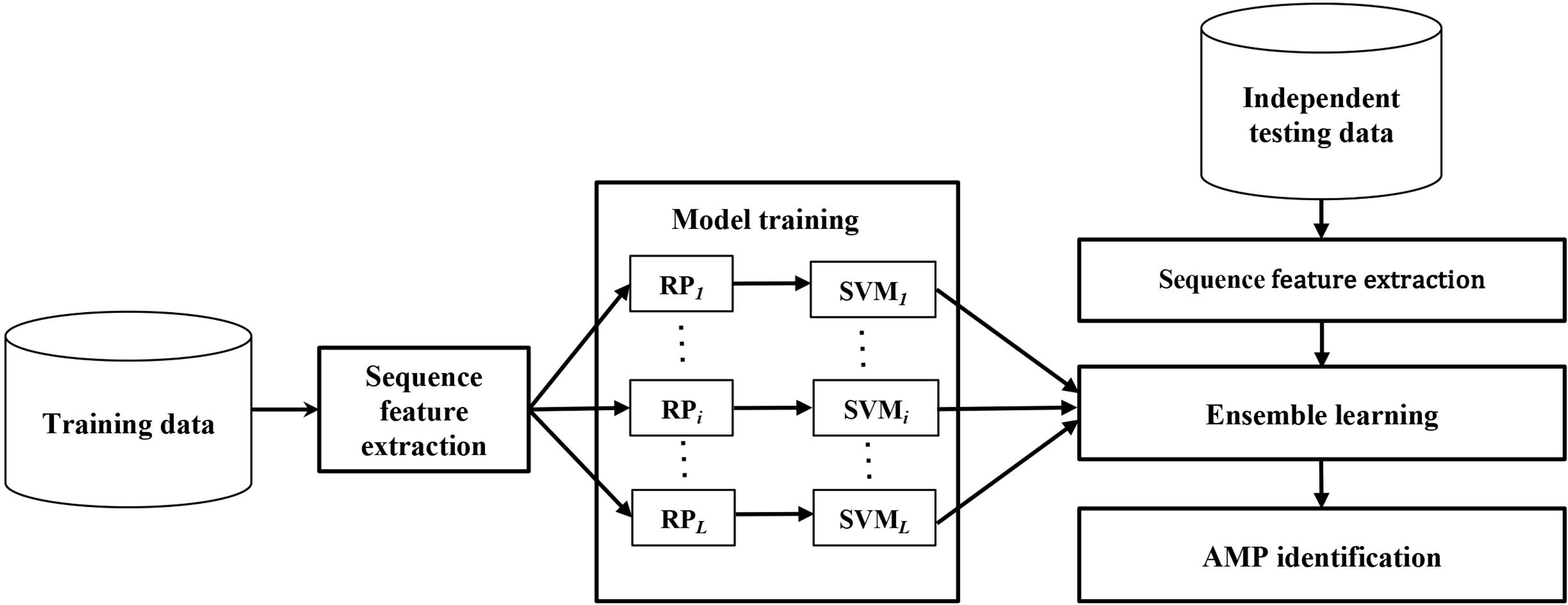
Schematic representation of SAMP workflow. Benchmarking data consisting of AMPs and non-AMPs were used for training. Features including our proposed proportionalized split amino acid composition (PSAAC) as well as conventional sequence features were constructed. Random projection (RP) was applied multiple times to reduce the feature dimension for robustness. For each RP, the feature matrix was transformed in a low-dimensional space and was then fed into a classification model (here we used a radial basis function (RBF) based support vector machine (SVM) model). The decision scores generated by the RBF-SVM model were integrated by an ensemble learning scheme, based on which predictions for independent test data were made to identify AMPs.

We then compared and selected the appropriate classifier for AMP sequences classification, including RF, LR, SVM, multilayer perceptron (MLP) and XGBoost. Specifically, SVM is a widely used classification model that allows for the use of different kernel functions to make predictions on both linear and non-linear data. The model is characterized by several parameters, including the regularization parameter (C), the choice of kernel function, and the kernel’s gamma parameter (such as Radial Basis Function (RBF)). Random Forest is a powerful ensemble learning method used for both classification and regression tasks. It works by building multiple decision trees and merging their outputs to make predictions. Hyperparameters like n_estimators, max_depth, min_samples_split, min_samples_leaf and bootstrap need to be optimized. Logistic regression makes predictions by modeling the relationship between variables based on logistic/sigmoid function, which is a recognized powerful algorithm used for binary variable classification. Its hyperparameter contains regularization parameter (C), penalty type and solver type. MLP consists at least three layers of notes, including input layer, one/more hidden layers, and an output layer, and it has been used widely for classification and regression analysis. All notes except the input, apply a nonlinear activation function and utilize backpropagation for training. XGBoost is designed for gradient boosting specifically with high performance and scalability, based on the combination of multiple decision trees to create a strong prediction model. It includes the parameter of max_depth, learning_rate and n_estimators. In each iteration of five classifier models, we trained them by performing grid search with repeated 10-fold cross validation to search for the best hyperparameters. Then, the model with the best hyperparameters was used to generate decision function scores for the independent testing datasets. Subsequently, based on the prediction performance, the classifier demonstrating the highest accuracy will be selected to form the foundational architecture of SAMP.

### Overview of SAMP

SAMP is an ensemble-based model that accurately classifies antimicrobial peptides by averaging the prediction scores from a set of base SVM models. Importantly, SAMP introduces the PSAAC feature, in addition to the widely used numeric features for antimicrobial peptide prediction task proposed in [53]. By implementing the ensemble technique and including a novel feature set, SAMP can excel performance of state-of-the-art approaches.

SAMP first encoded the peptide sequence into numeric features, such as AAC, PHYC, and PSAAC (**Fig. 2**). The features were then scaled and projected to a pre-defined lower dimension using random projection technique. Base SVM models were built to generate the prediction scores for each run, which were eventually integrated by an ensemble learning scheme. SAMP was then evaluated on independent test data from four species (including amphibian, bacteria, human, and plant) and compared to other state-of-the-art methods, including iAMPpred and AMPScanner V2. To make fair comparisons, the same training data and independent test data were used to compare SAMP and other state-of-the-art methods.

Overall, the PSAAC enables SAMP to capture the peptide sequence information from both the middle region and the N/C-terminus regions, which significantly boosts the model performance in comparison to state-of-the-art methods. In the following sections, we demonstrate the superb performance of SAMP across datasets from different species.

### Benchmarking with the state-of-the-art methods

We compared the performance of our model with two state-of-the-art methods, iAMPpred and AMPScanner V2. The benchmark test was performed by using the AMP and non-AMP data collected from the dbAMP database. The training data reported in the papers [50,53] for iAMPpred and AMPScanner V2 were obtained to train SAMP separately. To demonstrate the importance of our proposed feature PSAAC and the robustness of our ensemble based SVM model design, we conducted two types of further analyses. First, we trained models both with and without the PSAAC features, evaluating the results to ascertain the importance of PSAAC. Following this, we employed both the ensemble based SVM model design and basic SVM model with one time RP for training and assessed their respective performances. For performance evaluation, we considered four major metrics: accuracy, MCC, G-measure and F1-score. Here, MCC is a measure which produces high score only if the prediction obtained good performance in all four aspects, true and false positives and negatives, of the confusion matrix, making it a reliable rate particularly for imbalanced datasets, as it is not biased toward the majority class [68]. The closer the value of MCC is to 1, the better the prediction effect of the classifier is. G-measure is the geometric mean of precision and recall, where the precision is the number of true positive cases divided by the number of all predicted as positive samples, and the recall is the number of true positive results divided by the number of all samples which should be regarded as positive. G-measure effectively balances the extreme ratio of positive to negative instances and the value ranges from 0 to 1, then a value closer to 1, indicating the classifier is performing well in both predicting the positive cases and maintaining accuracy, conversely, a value closer to 0 indicates bad performance. F1-score is the harmonic mean of precision and recall, equally weighting the two values. It differs from G-measure in that, F1-score is more sensitive to the extreme values, if there is low precision or recall, the F1-score decreases significantly, however, g-measure will be more tolerant. Similarly, a closer value to 1 means the better prediction ability of the classifier.

## Results

### Model performance and classifier selection

To enhance the prediction capability of SAMP, we initially selected five ML classifiers, namely SVM, RF, LR, MLP and XGBoost, using the same training and independent test dataset to train and test, then evaluated their performance. We performed 10-fold cross validation for 10 times, each time will get an assessment value, as shown in **Fig. 4**, SVM had better performance than LR, MLP, RF and XGBoost, and LR always presents the worst result, based on accuracy, MCC, G-measure and F1-score. Then, five trained classifiers were applied to predict labels for independent test data, as shown in **Fig. 5**, SVM exhibited the highest accuracy, MCC, G-measure and F1-score among all four test datasets. In summary, SVM presents a better performance than RF, MLP, XGBoost and LR, which was determined to serve as the basement of SAMP for further analysis.

**Fig. 4.**
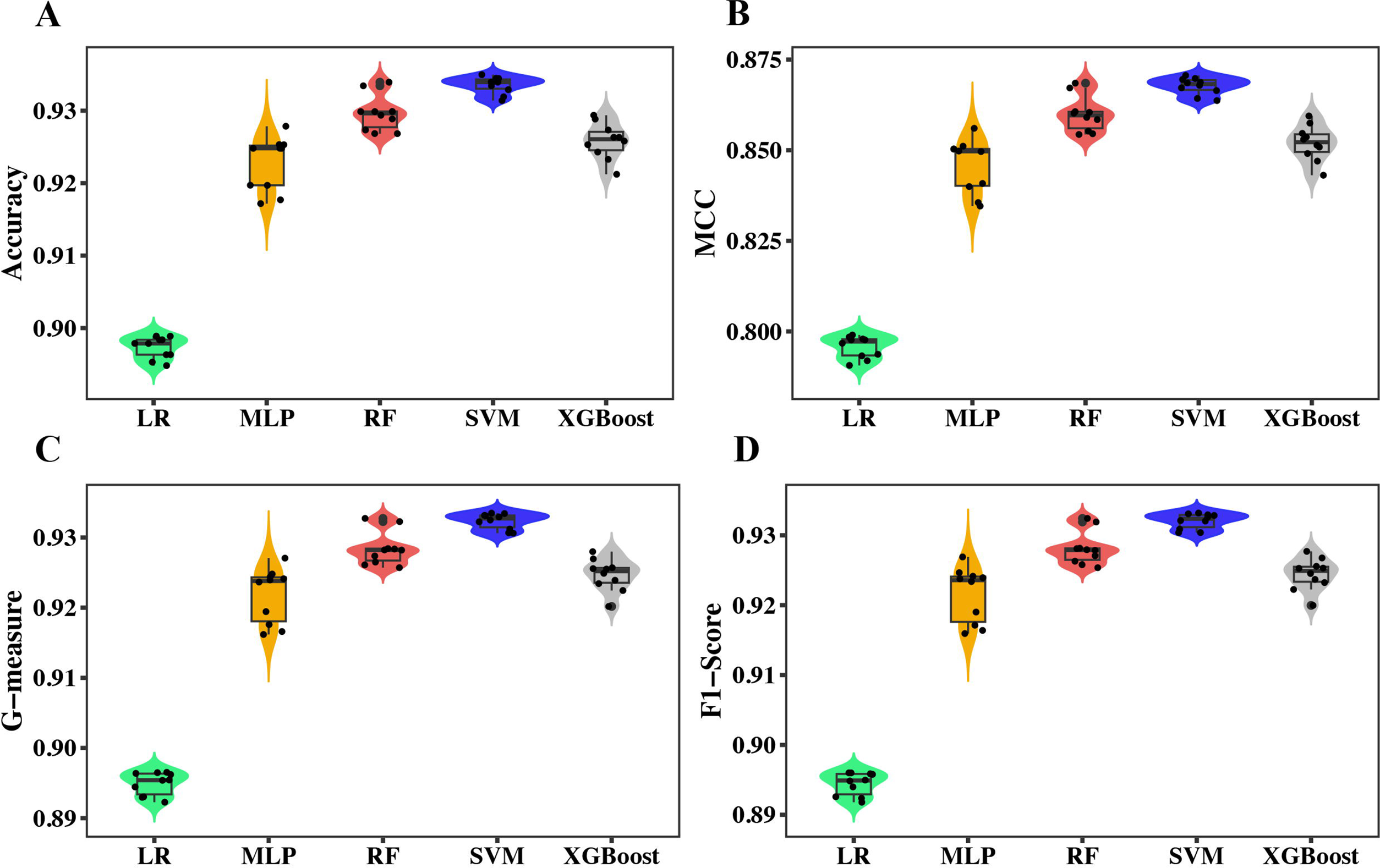
Comparing different classifiers for SAMP. All classifiers were trained on the same dataset collected from iAMPpred to perform 10 times of 10-fold cross validations. Performance measures based on (**A**) Accuracy, (**B**) MCC, (**C**) G-measure, and (**D**) F1-score were reported. Classifiers include logistic regression (LR), deep learning like multi-layer perceptron (MLP), random forest (RF), SVM (support vector machine), and XGBoost.

**Fig. 5.**
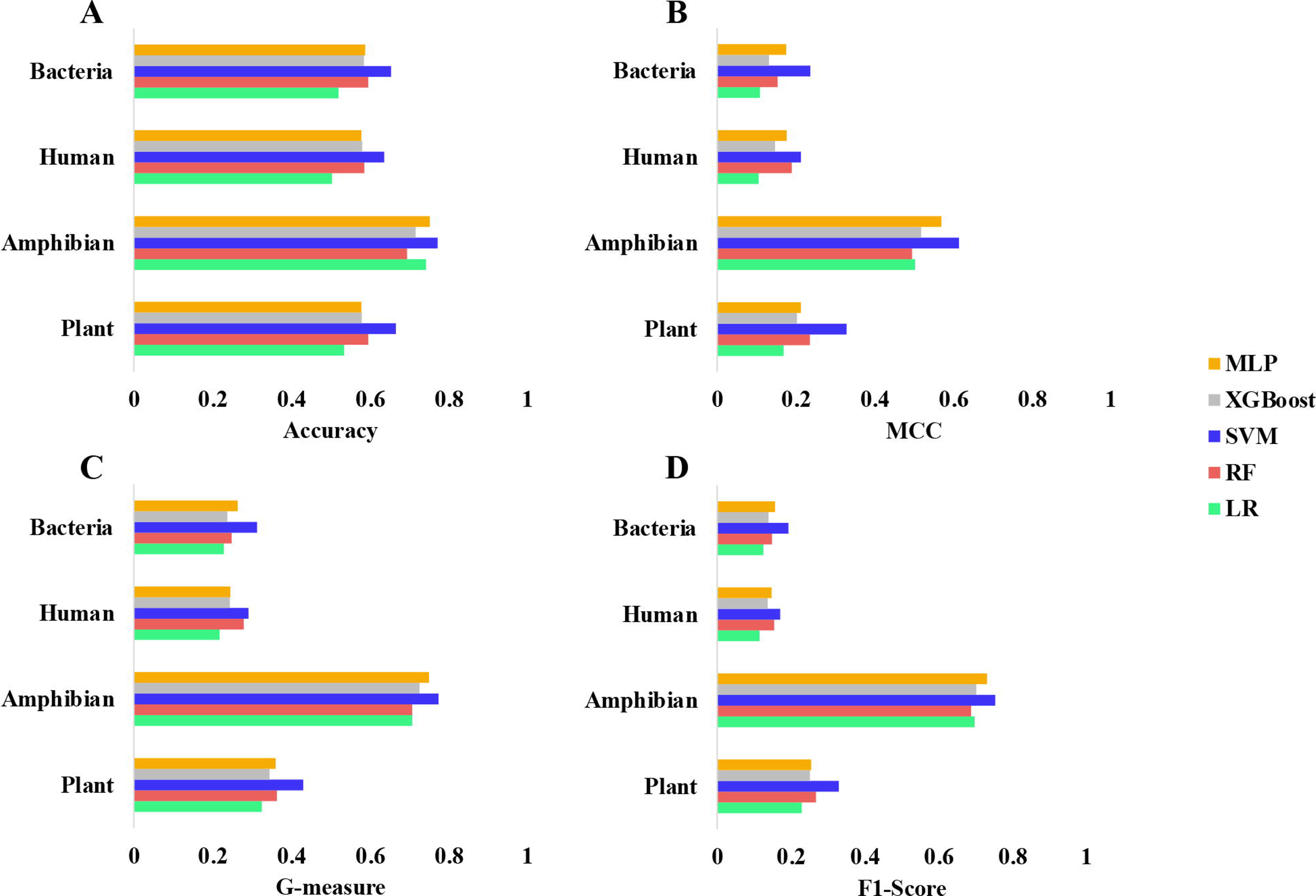
Comparison five machine learning models based on independent tests across multiple species. (**A**) Accuracy, (**B**) MCC, (**C**) G-measure, and (**D**) F1-score were compared across all species including bacteria, human, amphibian, and plant. All models were trained on the dataset collected from [53] and tested on independent test datasets collected from [60].

We also measured the performance of SAMP across different dimensions of random projection and all the possible proportions of PSAAC (**Table 1**). Specifically, we trained SAMP on the data collected from iAMPpred and AMPScanner V2 respectively. Grid-search with repeated 10-fold cross validation was applied to assess the model performance on training datasets. The number of dimensions used in RP was 50, 100, and 150. Importantly, the novel feature PSAAC enables a customized proportion of information to be obtained from a peptide sequence. To this end, we also evaluated the effect of different proportions of PSAAC on model performance. A given peptide sequence was first split into three parts according to the proportions specified. Next, the amino acid composition within each split was calculated, resulting in a total of 60 new features (see Method). The proportions evaluated include 2:2:6, 6:2:2, 2:6:2, and 3:4:3, where, for example, 2:2:6 represents cutting the peptide sequence from the N-terminus for 20% of the total sequence length, another 20% in the middle, and the remaining 60% for the C-terminus.

**Table 1.** Comparing different splitting schemes and reduced dimensions for SAMP. The splitting scheme means different ratios of the sequence lengths of the N-terminus region, the middle region, and C-terminus region. For example, 2:2:6 means splitting a peptide into three regions as the N-terminus region accounting for 20% of the total sequences, the middle region 20%, and the C-terminus region 60%. Here we tried four different splitting schemes including 2:2:6, 6:2:2, 2:6:2, and 3:4:3. For reduced dimensions of features, we tried three different cases, 50, 100, and 150. ACC, accuracy; MCC, Matthews correlation coefficient; Sn, sensitivity; Sp, specificity; AUC, area under the receiver operating characteristic curve. Numbers in bold represent the best performance for each splitting scheme.

| PSAAC | Dimensions | Evaluation Metrics |  |  |  |  |
| --- | --- | --- | --- | --- | --- | --- |
| Split |  | ACC | MCC | Sn | Sp | AUC |
| 2:2:6 | 50 | 93.04 | 86.04 | 91.06 | 94.92 | 97.58 |
|  | 100 | **93.29** | 86.24 | 91.16 | **95.02** | **97.79** |
|  | 150 | 93.09 | **86.43** | **91.57** | 94.82 | 97.77 |
| 6:2:2 | 50 | 93.24 | 86.28 | 90.65 | 95.53 | 97.63 |
|  | 100 | 93.24 | **86.70** | **90.75** | **95.83** | 97.68 |
|  | 150 | **93.29** | 86.59 | **90.75** | 95.73 | **97.72** |
| 2:6:2 | 50 | 93.09 | 86.29 | 90.55 | **95.63** | 97.63 |
|  | 100 | 93.60 | 87.24 | **91.97** | 95.22 | **97.87** |
|  | 150 | **93.65** | **87.35** | **91.97** | 95.33 | 97.82 |
| 3:4:3 | 50 | **93.65** | **87.36** | 91.77 | **95.53** | 97.83 |
|  | 100 | 93.39 | 86.85 | 91.57 | 95.22 | 97.89 |
|  | 150 | **93.65** | 87.34 | **92.07** | 95.22 | **97.93** |

As shown in **Table 1**, it presented a comprehensive overview of the SAMP performance under varying ratios of PSAAC with different dimensions. It emphasized how different splitting schemes influenced the performance of SAMP, such as ACC, MCC, Sn, Sp, and AUC. The ACC presented minimum variation, ranging from 93.04 to 93.65 which indicated a consistently good performance across different configurations. The MCC, Sn, Sp and AUC values varied slightly more but still could demonstrate the robust performance of SAMP, with MCC ranging from 86.06 to 87.36, Sn from 90.55 to 92.07, Sp from 94.82 to 95.83, and AUC ranging from 97.58 to 97.93. Among all the configurations, the 6:2:2 PSAAC ratio reached the highest Sp, while the 2:6:2 ratio got the best accuracy and the 3:4:3 ratio outperformed others in terms of ACC, MCC, Sn, and AUC. Analyzing performance based on dimensions, obviously, the dimension of 50 led in ACC and MCC, the dimension of 100 exceeding in Sp, and the dimension of 150 topped in ACC, Sn, and AUC. Therefore, the best model performance was achieved when the proportion was 2:6:2 with feature dimensions reduced to 150 using RP, which indicated the importance of the peptide sequence information from the middle region of peptides.

### Benchmarking with the state-of-the-art methods

To further evaluate the predictive performance of SAMP, we compared it with the performance of two state-of-the-art AMP prediction tools, iAMPpred and AMPScanner V2. We first retrained SAMP with the same training data from the two methods respectively. We compared their performance by using datasets collected from the dbAMP database. In particular, we chose the AMPs and non-AMPs from plants, bacteria, amphibians, and humans. We considered accuracy, MCC, G-measure and F-1 score as our major evaluation metrics.

First, SAMP was trained on 984 AMPs and 984 non-AMPs obtained from the iAMPpred paper. The trained SAMP was tested on the independent dataset from dbAMP. To assess the performance of iAMPpred, we uploaded the independent testing dataset to their web portal (http://cabgrid.res.in:8080/amppred/). Similarly, we trained SAMP using the exact same training dataset from AMPScanner V2 and uploaded the testing data to the web portal provided on https://www.dveltri.com/ascan/v2/ascan.html. As shown in **Fig. 6**, SAMP demonstrates better performance compared to both iAMPpred and AMPScanner across all four metrics: accuracy, MCC, G-measure and F1-score. When specifically comparing SAMP with iAMPpred (**Fig. 6A**), the most obvious advantage of SAMP is observed in MCC for predicting amphibian labels, where SAMP is 73% more accurate than iAMPpred. On the other hand, the smallest difference is noticed in the F1-score for predicting human labels, with SAMP being 11% more effective than iAMPpred. Notably, all MCC values for iAMPpred are negative, indicating this tool may predict adverse results. Comparing SAMP with AMPScanner (**Fig. 6B**) reveals similar trends. Probably due to a smaller data set in the APD, the largest disparity is seen in Accuracy for human AMP predictions, where SAMP shows a 29% improvement over AMPScanner, whereas the smallest difference is in the G-measure for human predictions, with a small improvement of 8% by SAMP over AMPScanner.

**Fig. 6.**
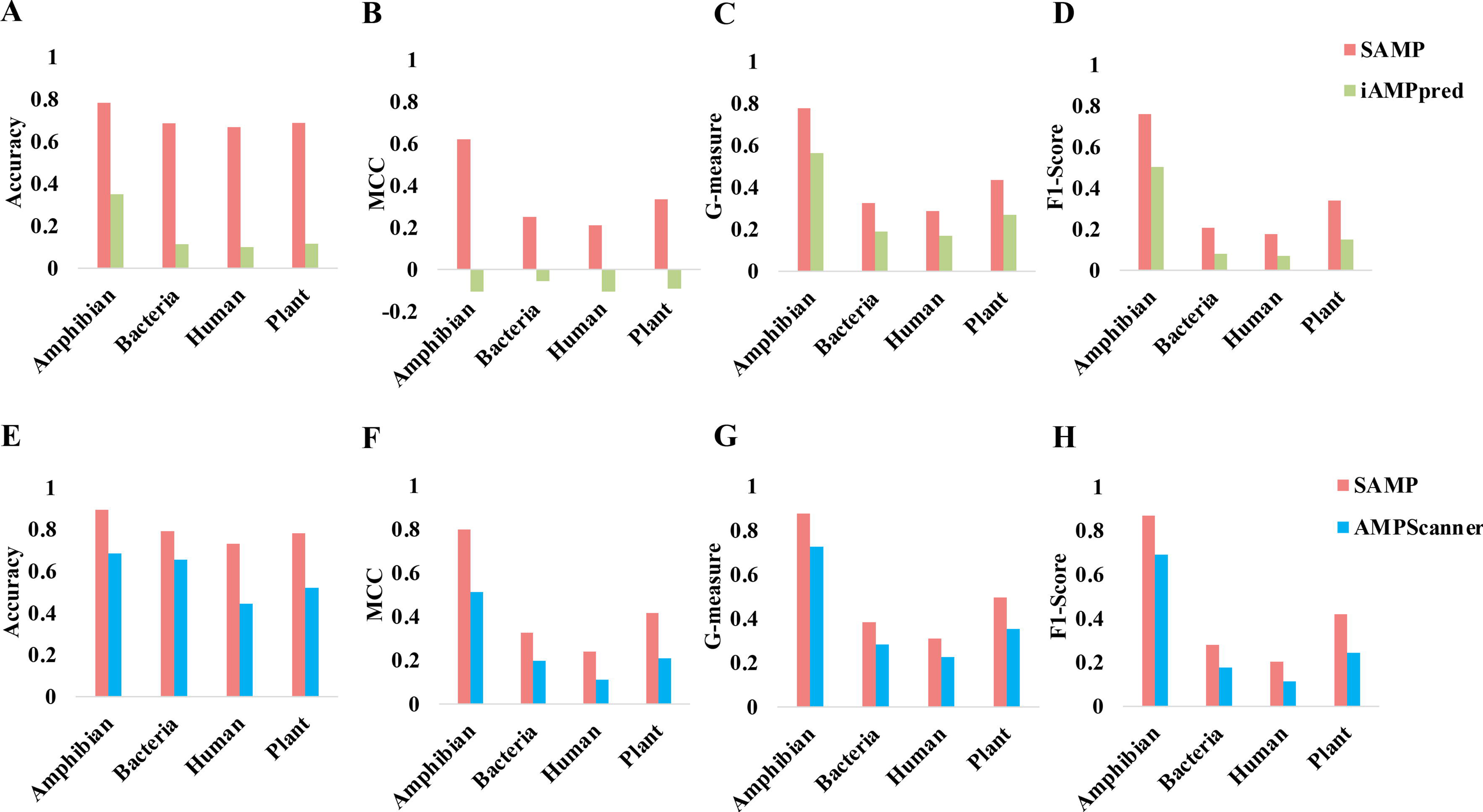
Comparing SAMP with state-of-the-art methods on different species datasets. Comparing SAMP and iAMPpred across different species in terms of (**A**) Accuracy, (**B**) MCC, (**C**) G-measure and (**D**) F1-score. SAMP was trained on the same training dataset collected from iAMPpred and tested on independent test dataset collected from dbAMP. Comparing SAMP and AMPScanner V2 across different species in terms of (**E**) Accuracy, (**F**) MCC, (**G**) G-measure and (**H**) F1-score. SAMP was trained on the same training dataset collected from AMPScanner V2 and tested on independent test dataset collected from dbAMP.

Furthermore, we evaluated the impact of proportionalized split amino acid composition (PSAAC) and the ensemble-based SVM model architecture on the predictive performance (**Fig. 7**). After training with data from iAMPpred, SAMP consistently outperformed both the SAMP without the PSAAC feature and the vanilla SVM model without ensemble learning. This improvement was consistent in all label predictions. Specifically, SAMP demonstrated at least a 11% increase in accuracy, 9% in MCC, 5% in G-measure, and 7% in F1-score compared to the situation of deleting the PSAAC feature, and at least a 2% increase in accuracy, 1% in MCC, 1% in G-measure, 1% in F1-score compared to the situation of deleting the layer of ensemble learning. Similar outcomes were observed when trained with AMPScanner data, with SAMP outperforming the aforementioned situations across all measures.

**Fig. 7.**
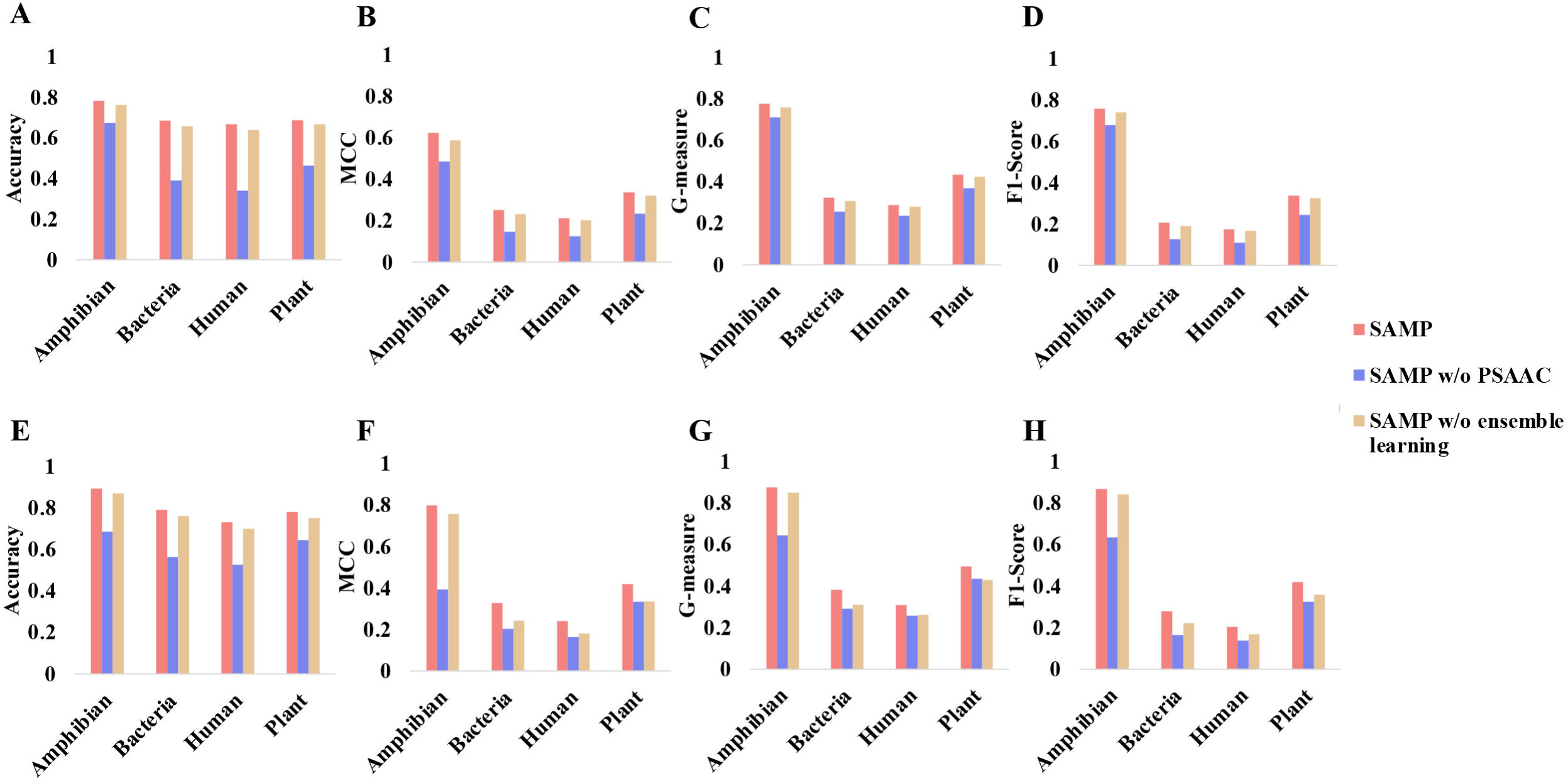
PSAAC and ensemble learning contribute to improving prediction performance of SAMP for identifying AMPs. Comparing SAMP and SAMP without the PSAAC feature across different species in terms of (**A**) Accuracy, (**B**) MCC, (**C**) G-measure and (**D**) F1-score. All models were trained on the same training dataset collected from iAMPpred and tested on independent test dataset collected from dbAMP. Comparing SAMP and SAMP without ensemble learning across different species in terms of (**E**) Accuracy, (**F**) MCC, (**G**) G-measure and (**H**) F1-score. All models were trained on the same training dataset collected from AMPScanner V2 and tested on independent test dataset collected from dbAMP.

### Feature scaling

A crucial step in improving the performance of SVM-based models is feature scaling. Intuitively, if the features are measured in different scales, the decision boundary calculation of SVM would be dominated by the features with the largest scales. In our study, we always scaled the features after the feature generation stage using the *scale* function in R. In particular, the peptide sequence features are calculated in different scales. For example, the amino acid composition is measured as some values between 0 and 1, but certain physio-chemical properties such as hydrophobicity can have various ranges of value. We believe this step is essential for SAMP to make accurate predictions and is worth experimenting. We generated two sets of features from the peptide sequences used to train iAMPpred, in which one set of features was scaled and the other was not. Two separate SAMP models were trained and evaluated on the independent test datasets. Our results indicate that scaling is indeed extremely important for SAMP, consistently boosting the model performance by a least 50% across datasets (**Table 2**).

**Table 2.** Scaling the features is crucial for SAMP for identifying AMPs. The scaling is performed by subtracting the mean of each feature and dividing by the feature’s standard deviation. Scaling is a crucial step for SAMP. ACC, accuracy; MCC, Matthews correlation coefficient; AUC, area under the receiver operating characteristic curve.

| Dataset | Metric | SAMP (Scaled) | SAMP (No Scale) |
| --- | --- | --- | --- |
| dbAMP Plant | Accuracy | 0.668 | 0.102 |
|  | AUC | 0.744 | 0.112 |
|  | MCC | 0.332 | -0.184 |
| dbAMP Bacteria | Accuracy | 0.647 | 0.071 |
|  | AUC | 0.703 | 0.088 |
|  | MCC | 0.234 | -0.165 |
| dbAMP Amphibian | Accuracy | 0.779 | 0.336 |
|  | AUC | 0.844 | 0.039 |
|  | MCC | 0.624 | -0.169 |
| dbAMP Human | Accuracy | 0.637 | 0.058 |
|  | AUC | 0.712 | 0.137 |
|  | MCC | 0.204 | -0.2 |

## Discussion

AMPs have gained greater attention as an alternative to chemical antibiotics. Indeed, some are already in applications either as antibiotics or as food preservatives [69]. Computational methods are developed as a supplement for wet lab experiments to design and identify AMPs, which reduces the cost and resources required. In this study, we present a novel ensemble-based model that achieves better AMP prediction performance than existing, state-of-the-art methods. To the best of our knowledge, SAMP is the first method that adopts PSAAC as one of the numeric features for AMP prediction tasks. Amino acid compositional splitting sheds new light on amino acid compositions of natural AMPs, which was initially discovered in 2009 [70]. In natural AMPs, alanine, glycine, leucine, and lysine are frequently occurring (or abundant) amino acids, while histidine, methionine, and tryptophan are least abundant amino acids. Our sequence splitting here reveals that leucine is preferentially dominant at the N-terminus of AMPs, while alanine is mainly located at the middle region. Glycine can appear frequently both at the N and the middle regions. In contrast, lysine is primarily abundant in the middle and C-terminus of natural AMPs. Interestingly after sequence splitting, the least abundant methionine and tryptophan appear mainly at the middle and the C-terminus regions, whereas histidine occupies the N-terminus. Also of note is that acidic glutamic acid is located at the N-terminus and acidic aspartic acid prefers the C-terminus region.

By combining this novel sequence-splitting feature with an ensemble-based SVM model architecture, SAMP is able to maximally extract peptide sequence information and outperform methods that apply either deep learning or traditional machine learning techniques. Additionally, we developed SAMP based on RP, a powerful dimension-reduction algorithm based on the Johnson–Lindenstrauss lemma [66] which can preserve the distances between data points while reducing the dimension [71]. As the number of data points continues to grow, the accuracy of prediction may be influenced due to the low efficiency of computational efficiency. Therefore, RP based models should have better performance compared to those without it. This has been evidenced in a large-scale single-cell RNA-sequencing (scRNA-seq) data processed algorithm which showed a higher classification efficiency under the contribution of ensemble RP layer [72]. As expected, our model with the ensemble RP layer also has a better performance as shown in **Fig. 7**. Our prediction also implies that data size influences prediction performance since the human AMPs, with the least data (<150 AMPs in the current APD), behave poorest compared to AMPs from bacteria, plants, and animals with more known positive data.

We also assessed the performance of SAMP with specific tools, like iAMPpred and AMPScanner V2, which are also designed for AMP prediction based on SVM and DNN respectively. SAMP proved slightly better performance than AMPScanner V2 and obviously higher accuracy than iAMPpred. Possible explanation for this discrepancy should be the omission of PSAAC and ensemble RP layer. Overall, this newly designed tool, SAMP, is expected to compensate for the existing tools for AMP prediction.

For future research directions, we will consider different ensemble methods by including more diverse model categories to improve the prediction accuracy. With the advance of deep learning, it would be appealing to investigate the performance of DL based models combined with PSAAC features, or whether the deep neural networks are able to capture the PSAAC features within their embedding space.

### Key points

We propose a novel method called SAMP that develops a new type of features called proportionalized split amino acid composition (PSAAC) to significantly boost the performance of identifying antimicrobial peptides.

PSAAC can identify residue patterns at both the N-terminus and the C-terminus as well as to retain sequence order information from the middle region of peptide fragments.

SAMP leverages an ensemble learning framework based on random projection to integrate various classifiers into a cohesive framework, effectively improving the performance accuracy.

SAMP outperforms state-of-the-art methods for AMP identification in terms of accuracy, G-measure, MCC and F1-score.

SAMP is a versatile tool capable of identifying AMPs from a variety of organisms including human, plant, bacteria and amphibian.

## Supplementary Data

**Supplementary Fig. S1.** Amino acid distribution in AMPs and non-AMPs datasets based on the dataset collected from AMPScanner V2. Amino acid distribution of all sequences in (**A**) AMPs and (**B**) non-AMPs dataset. Distribution of amino acid sequences in the N-terminus region, the middle region and the C-terminus region of (**C**) AMPs Dataset and (**D**) non-AMPs dataset.

**Supplementary Fig. S2.** Amino acid distribution in AMPs and non-AMPs datasets based on the dataset collected from dbAMP. Amino acid distribution of all sequences in (**A**) AMPs and (**B**) non-AMPs dataset. Distribution of amino acid sequences in the N-terminus region, the middle region and the C-terminus region of (**C**) AMPs Dataset and (**D**) non-AMPs dataset.

## Authors’ contributions

SW conceived and designed the study. JF and MS developed the algorithm, performed the experiments and analyzed the data. JF implemented the SAMP package. All authors participated in writing the paper. The manuscript was approved by all authors.

## Data availability

All the data used in this manuscript are publicly available in the corresponding references.

## Reference

1. Mishra NN, Bayer AS. Correlation of Cell Membrane Lipid Profiles with Daptomycin Resistance in Methicillin-Resistant Staphylococcus aureus. Antimicrob Agents Chemother 2013; 57:1082–1085

2. Fernandes P. Antibacterial discovery and development—the failure of success? Nature biotechnology 2006; 24:1497–1503

3. Adedeji WA. THE TREASURE CALLED ANTIBIOTICS. Ann Ib Postgrad Med 2016; 14:56–57

4. Thomas L. The youngest science: notes of a medicine-watcher. 1995;

5. Aminov RI. A brief history of the antibiotic era: lessons learned and challenges for the future. Frontiers in microbiology 2010; 1:134

6. Hutchings MI, Truman AW, Wilkinson B. Antibiotics: past, present and future. Current opinion in microbiology 2019; 51:72–80

7. Prestinaci F, Pezzotti P, Pantosti A. Antimicrobial resistance: a global multifaceted phenomenon. Pathogens and global health 2015; 109:309–318

8. De Oliveira DM, Forde BM, Kidd TJ, et al. Antimicrobial resistance in ESKAPE pathogens. Clinical microbiology reviews 2020; 33:10.1128/cmr.00181-19

9. Huemer M, Mairpady Shambat S, Brugger SD, et al. Antibiotic resistance and persistence— Implications for human health and treatment perspectives. EMBO reports 2020; 21:e51034

10. Frieri M, Kumar K, Boutin A. Antibiotic resistance. Journal of infection and public health 2017; 10:369–378

11. Lei J, Sun L, Huang S, et al. The antimicrobial peptides and their potential clinical applications. American journal of translational research 2019; 11:3919

12. Kadri SS. Key takeaways from the US CDC’s 2019 antibiotic resistance threats report for frontline providers. Critical care medicine 2020; 48:939–945

13. de Kraker MEA, Stewardson AJ, Harbarth S. Will 10 Million People Die a Year due to Antimicrobial Resistance by 2050? PLoS Med 2016; 13:e1002184

14. O’Neill J. Tackling drug-resistant infections globally: final report and recommendations. 2016;

15. Kang X, Dong F, Shi C, et al. DRAMP 2.0, an updated data repository of antimicrobial peptides. Scientific data 2019; 6:148

16. Chen CH, Lu TK. Development and challenges of antimicrobial peptides for therapeutic applications. Antibiotics 2020; 9:24

17. Mookherjee N, Anderson MA, Haagsman HP, et al. Antimicrobial host defence peptides: functions and clinical potential. Nature reviews Drug discovery 2020; 19:311–332

18. Diamond G, Beckloff N, Weinberg A, et al. The roles of antimicrobial peptides in innate host defense. Current pharmaceutical design 2009; 15:2377–2392

19. Wang G, Li X, Wang Z. APD3: the antimicrobial peptide database as a tool for research and education. Nucleic Acids Res 2016; 44:D1087–D1093

20. Hiemstra PS, Amatngalim GD, Van Der Does AM, et al. Antimicrobial Peptides and Innate Lung Defenses. Chest 2016; 149:545–551

21. Silva ON, De La Fuente-Núñez C, Haney EF, et al. An anti-infective synthetic peptide with dual antimicrobial and immunomodulatory activities. Scientific reports 2016; 6:35465

22. Frohm M, Agerberth B, Ahangari G, et al. The expression of the gene coding for the antibacterial peptide LL-37 is induced in human keratinocytes during inflammatory disorders. Journal of Biological Chemistry 1997; 272:15258–15263

23. Liang W, Diana J. The dual role of antimicrobial peptides in autoimmunity. Frontiers in immunology 2020; 11:545577

24. De La Fuente-Núñez C, Silva ON, Lu TK, et al. Antimicrobial peptides: Role in human disease and potential as immunotherapies. Pharmacology & Therapeutics 2017; 178:132–140

25. Li C, Zhu C, Ren B, et al. Two optimized antimicrobial peptides with therapeutic potential for clinical antibiotic-resistant Staphylococcus aureus. European journal of medicinal chemistry 2019; 183:111686

26. Fan L, Wei Y, Chen Y, et al. Epinecidin-1, a marine antifungal peptide, inhibits Botrytis cinerea and delays gray mold in postharvest peaches. Food Chemistry 2023; 403:134419

27. Adade CM, Oliveira IR, Pais JA, et al. Melittin peptide kills Trypanosoma cruzi parasites by inducing different cell death pathways. Toxicon 2013; 69:227–239

28. Huan Y, Kong Q, Mou H, et al. Antimicrobial peptides: classification, design, application and research progress in multiple fields. Frontiers in microbiology 2020; 11:582779

29. Wachinger M, Kleinschmidt A, Winder D, et al. Antimicrobial peptides melittin and cecropin inhibit replication of human immunodeficiency virus 1 by suppressing viral gene expression. Journal of General Virology 1998; 79:731–740

30. Torres MD, Cao J, Franco OL, et al. Synthetic biology and computer-based frameworks for antimicrobial peptide discovery. ACS nano 2021; 15:2143–2164

31. Li J, Koh J-J, Liu S, et al. Membrane active antimicrobial peptides: translating mechanistic insights to design. Frontiers in neuroscience 2017; 11:244777

32. Travkova OG, Moehwald H, Brezesinski G. The interaction of antimicrobial peptides with membranes. Advances in colloid and interface science 2017; 247:521–532

33. Kumar P, Kizhakkedathu JN, Straus SK. Antimicrobial peptides: diversity, mechanism of action and strategies to improve the activity and biocompatibility in vivo. Biomolecules 2018; 8:4

34. Ahmed TA, Hammami R. Recent insights into structure–function relationships of antimicrobial peptides. Journal of food biochemistry 2019; 43:e12546

35. Le C-F, Fang C-M, Sekaran SD. Intracellular targeting mechanisms by antimicrobial peptides. Antimicrobial agents and chemotherapy 2017; 61:10.1128/aac.02340-16

36. Li S, Wang Y, Xue Z, et al. The structure-mechanism relationship and mode of actions of antimicrobial peptides: A review. Trends in Food Science & Technology 2021; 109:103–115

37. Andersson DI, Hughes D, Kubicek-Sutherland JZ. Mechanisms and consequences of bacterial resistance to antimicrobial peptides. Drug Resistance Updates 2016; 26:43–57

38. Lázár V, Martins A, Spohn R, et al. Antibiotic-resistant bacteria show widespread collateral sensitivity to antimicrobial peptides. Nature microbiology 2018; 3:718–731

39. Spohn R, Daruka L, Lázár V, et al. Integrated evolutionary analysis reveals antimicrobial peptides with limited resistance. Nature communications 2019; 10:4538

40. Cortes C, Vapnik V. Support-vector networks. Machine learning 1995; 20:273–297

41. Breiman L. Random forests. Machine learning 2001; 45:5–32

42. Tolles J, Meurer WJ. Logistic regression: relating patient characteristics to outcomes. Jama 2016; 316:533–534

43. Wang G, Vaisman II, Van Hoek ML. Machine Learning Prediction of Antimicrobial Peptides. Computational Peptide Science 2022; 2405:1–37

44. Huang J, Xu Y, Xue Y, et al. Identification of potent antimicrobial peptides via a machine-learning pipeline that mines the entire space of peptide sequences. Nature Biomedical Engineering 2023; 7:797–810

45. Chen T, Guestrin C. Xgboost: A scalable tree boosting system. Proceedings of the 22nd acm sigkdd international conference on knowledge discovery and data mining 2016; 785–794

46. LeCun Y, Bottou L, Bengio Y, et al. Gradient-based learning applied to document recognition. Proceedings of the IEEE 1998; 86:2278–2324

47. Hochreiter S, Schmidhuber J. Long short-term memory. Neural computation 1997; 9:1735– 1780

48. Ma Y, Guo Z, Xia B, et al. Identification of antimicrobial peptides from the human gut microbiome using deep learning. Nature Biotechnology 2022; 40:921–931

49. Devlin J, Chang M-W, Lee K, et al. Bert: Pre-training of deep bidirectional transformers for language understanding. arXiv preprint arXiv:1810.04805 2018;

50. Veltri D, Kamath U, Shehu A. Deep learning improves antimicrobial peptide recognition. Bioinformatics 2018; 34:2740–2747

51. Yan J, Bhadra P, Li A, et al. Deep-AmPEP30: improve short antimicrobial peptides prediction with deep learning. Molecular Therapy-Nucleic Acids 2020; 20:882–894

52. Lee KJ. Architecture of neural processing unit for deep neural networks. Advances in Computers 2021; 122:217–245

53. Meher PK, Sahu TK, Saini V, et al. Predicting antimicrobial peptides with improved accuracy by incorporating the compositional, physico-chemical and structural features into Chou’s general PseAAC. Sci Rep 2017; 7:42362

54. García-Jacas CR, Pinacho-Castellanos SA, García-González LA, et al. Do deep learning models make a difference in the identification of antimicrobial peptides? Briefings in Bioinformatics 2022; 23:bbac094

55. Bingham E, Mannila H. Random projection in dimensionality reduction: applications to image and text data. Proceedings of the seventh ACM SIGKDD international conference on Knowledge discovery and data mining 2001; 245–250

56. Wan S, Mak M-W, Kung S-Y. Ensemble linear neighborhood propagation for predicting subchloroplast localization of multi-location proteins. Journal of proteome research 2016; 15:4755–4762

57. Wang Z. APD: the Antimicrobial Peptide Database. Nucleic Acids Research 2004; 32:590D – 592

58. Wang G. The antimicrobial peptide database is 20 years old: Recent developments and future directions. Protein Science 2023; 32:e4778

59. Lata S, Sharma B, Raghava G. Analysis and prediction of antibacterial peptides. BMC Bioinformatics 2007; 8:263

60. Jhong J-H, Yao L, Pang Y, et al. dbAMP 2.0: updated resource for antimicrobial peptides with an enhanced scanning method for genomic and proteomic data. Nucleic acids research 2022; 50:D460–D470

61. Osorio D, Rondón-Villarreal P, Torres R. Peptides: a package for data mining of antimicrobial peptides. Small 2015; 12:44–444

62. Verma R, Varshney GC, Raghava GPS. Prediction of mitochondrial proteins of malaria parasite using split amino acid composition and PSSM profile. Amino Acids 2010; 39:101–110

63. Hayat M, Khan A, Yeasin M. Prediction of membrane proteins using split amino acid and ensemble classification. Amino acids 2012; 42:2447–2460

64. Nakai K. Protein sorting signals and prediction of subcellular localization. Advances in protein chemistry 2000; 54:277–344

65. Emanuelsson O. Predicting protein subcellular localisation from amino acid sequence information. Briefings in bioinformatics 2002; 3:361–376

66. Johnson WB, Lindenstrauss J, Schechtman G. Extensions of Lipschitz maps into Banach spaces. Israel Journal of Mathematics 1986; 54:129–138

67. Li P, Hastie TJ, Church KW. Very sparse random projections. Proceedings of the 12th ACM SIGKDD international conference on Knowledge discovery and data mining 2006; 287– 296

68. Chicco D, Jurman G. The advantages of the Matthews correlation coefficient (MCC) over F1 score and accuracy in binary classification evaluation. BMC genomics 2020; 21:1–13

69. Mishra B, Reiling S, Zarena D, et al. Host defense antimicrobial peptides as antibiotics: design and application strategies. Current opinion in chemical biology 2017; 38:87–96

70. Wang G, Li X, Wang Z. APD2: the updated antimicrobial peptide database and its application in peptide design. Nucleic Acids Research 2009; 37:D933–D937

71. Frankl P, Maehara H. The Johnson-Lindenstrauss lemma and the sphericity of some graphs. Journal of Combinatorial Theory, Series B 1988; 44:355–362

72. Wan S, Kim J, Won KJ. SHARP: hyperfast and accurate processing of single-cell RNA-seq data via ensemble random projection. Genome Res. 2020; 30:205–213

