## Supplementary figures and images for "SAMP: Identifying Antimicrobial Peptides by an Ensemble Learning Model Based on Proportionalized Split Amino Acid Composition"

### Supplementary Fig. S1

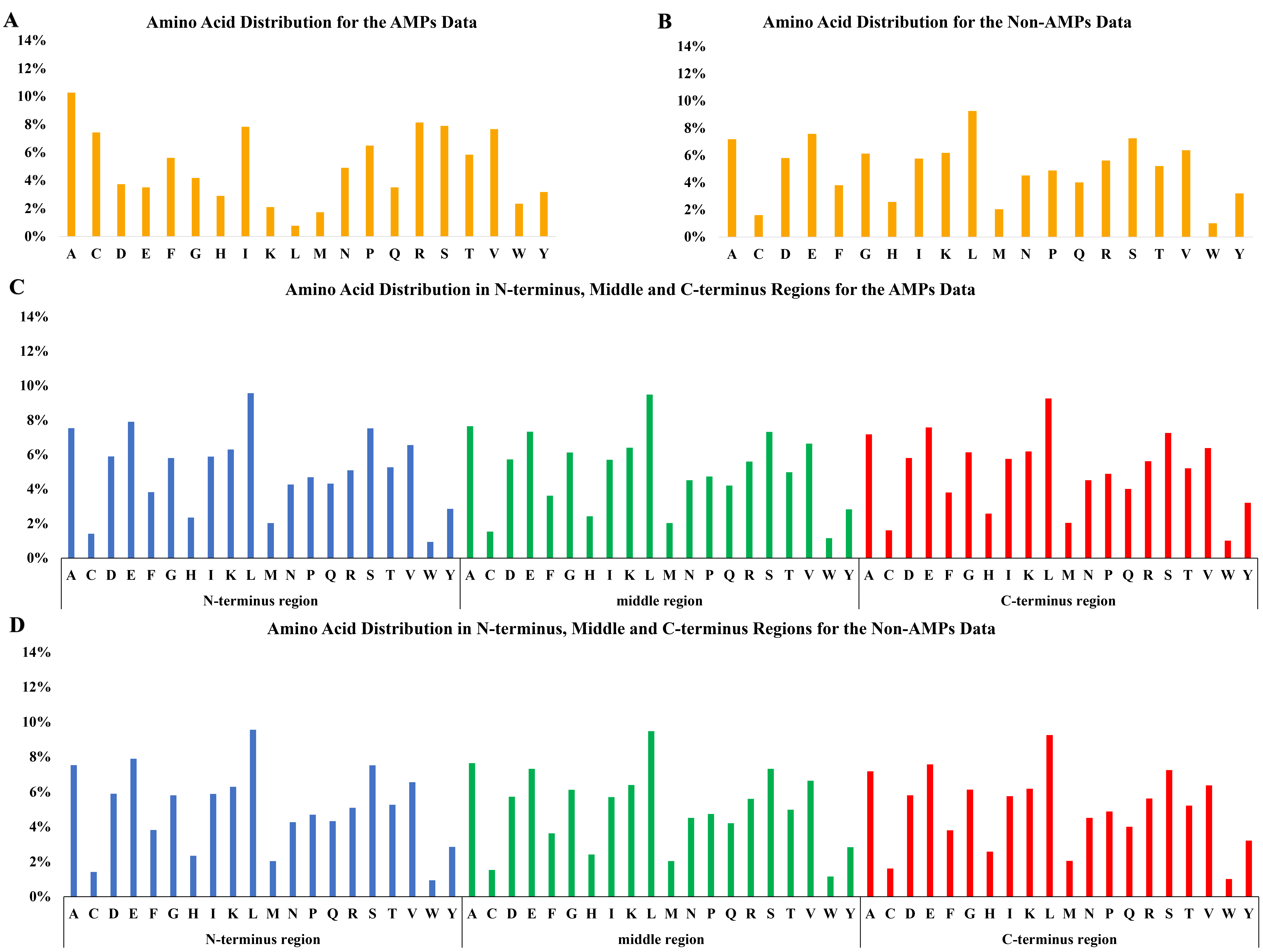

### Supplementary Fig. S2

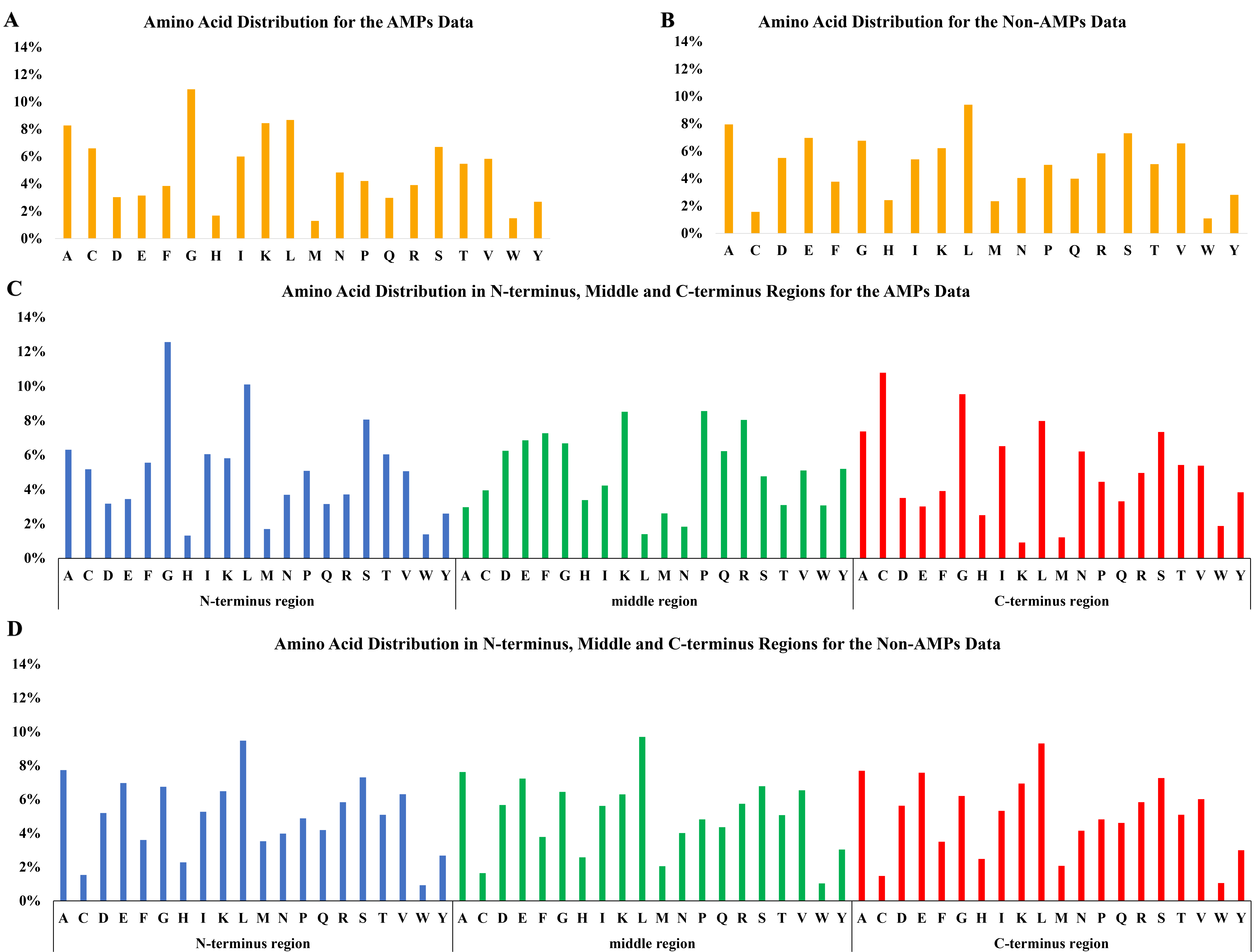
